# Single-cell RNA-seq of amniotic fluid provides unique insight into fetal development

**DOI:** 10.64898/2026.10.26.762078

**Authors:** Calandra V Grima, Ishara Atukorala, Sally Beard, Shalini Guleria, Liam Neil, Natalie Hannan, Bhupinder Pal, Jovana Maksimovic, Lisa Hui, Alicia Oshlack

**Affiliations:** Peter MacCallum Cancer Centre, Computational Biology Program, Parkville, Australia; The Sir Peter MacCallum Department of Oncology, University of Melbourne, Parkville, Australia; Department of Obstetrics and Gynaecology, University of Melbourne, Parkville, Australia; Department of Perinatal Medicine, Mercy Hospital for Women, Heidelberg, Australia; Murdoch Children’s Research Institute, Respiratory Group, Parkville, Australia; Olivia Newton-John Cancer Research Institute, Single Cell Cancer Genomics Laboratory, Heidelberg, Australia; Murdoch Children’s Research Institute, Reproductive Epidemiology Group, Parkville, Australia; School of Mathematics and Statistics, University of Melbourne, Melbourne, Victoria, Australia

## Abstract

**Background:** Human fetal development is difficult to study due to ethical and practical constraints on the collection and use of human fetal tissue. Amniotic fluid (AF) is routinely collected during clinical amniocentesis and contains live fetal cells shed from developing tissues. Thus, we use single-cell RNA-sequencing (scRNA-seq) to build the first cell atlas of amniotic fluid from healthy second-trimester fetuses.

**Methods:** scRNA-seq was performed on AF samples collected from seven euploid, structurally normal pregnancies at 15–25 weeks’ gestation, all of which resulted in live births of healthy infants. Mapping to a modified transcriptome was used to help reduce counts sparsity. Supervised cell type annotation, unsupervised clustering and marker gene analyses were used to determine the cell types present in AF. Gestational age-related differences between earlier (≤ 19 weeks) and later (≥ 23 weeks) gestation samples were examined using differential gene expression (DGE) and differential proportion analyses.

**Results:** Fetal skin cells were the most abundant cell type in AF, followed by myeloid and lymphoid cell types. We observed significant gestational age–related shifts in cell type composition, with an increased proportion of lung epithelium cells in later-age samples, coinciding with the canalicular phase of lung development. DGE analysis of fetal skin cells between earlier and later gestation revealed numerous gene expression changes that recapitulated known developmental processes across the second trimester. These were also enriched in an independent AF dataset from pregnancies with diverse fetal pathologies. Our dataset provides an annotated atlas of 11,414 healthy fetal human cells, permitting further study into normal second trimester development. We provide an integrated fetal cell atlas of 45,048 cells combining previous data with our healthy samples.

**Conclusions:** This study applied novel analytical approaches to demonstrate the utility of scRNA-seq in characterising AF and its capacity to capture a diverse range of viable fetal cell types. Our results suggest that changes in AF cellular composition mirror organ development, underscoring the potential of AF scRNA-seq to deepen our understanding of normal and pathological fetal biology.

## Introduction

Human fetal development is an important, yet challenging research field, as ethical and practical constraints on sample collection limit the study of live human fetal cells [1,2]. Despite these difficulties, much work has characterised the complex processes of fetal development, improving our understanding of many congenital conditions [2]. Fetal development research has great potential for shedding light on broader areas of human medicine, with growing evidence highlighting the role and reactivation of developmental processes in diseases such as cancer, as well as in furthering our understanding of regenerative medicine [2]. Additionally, the Developmental Origins of Health and Disease hypothesis (DOHaD) describes how all stages of development from pre-conception to infancy can impact the long-term health of an individual [3]. Thus, furthering our understanding of normal fetal development could help elucidate potential drivers of poorer adult health outcomes that originate during gestation [3].

A vital consideration in human fetal developmental research is the source of the samples, which are frequently derived from terminated/deceased pregnancies or from organoids [1,2]. The regulatory landscape of access to such tissues is complex worldwide, with region-specific limits on the ages and types of samples available [2]. These necessary guidelines therefore introduce a systematic bias in the samples that can be obtained for human fetal cell atlases, which are largely of early gestational age [4]. Thus, a crucial, yet unexplored avenue of human fetal development research is non-destructively examining samples from live ongoing pregnancies throughout gestation.

Amniotic fluid (AF) is a complex biofluid that surrounds the fetus in utero [5]. As AF is in constant contact with the developing fetus, it contains a suspension of viable cells and debris [5], as well as cell-free nucleic acids [6] and extracellular vesicles [7]. Thus, studying AF has the potential to provide insights into the development of multiple organ systems, including tissues not in direct contact with the amniotic cavity [8]. A key advantage of studying AF is its relative ease of access. Amniocentesis can be performed for prenatal diagnosis of fetal genetic conditions or congenital infections from 15 weeks onwards [9]. The AF transcriptome of normal pregnancies has been previously described using cell-free RNA in AF supernatant, providing insight into the cellular processes occurring in fetus at a given timepoint [6]. However, cell-free RNA by definition arises from cells that have lysed and describes a bulk transcriptional profile [6]. In contrast, single-cell RNA-sequencing (scRNA-seq) of AF allows the transcriptional profile of individual, live cells shed by a fetus to be examined [1]. Here, we present a fetal cell atlas of AF from healthy second-trimester pregnancies, all resulting in term or near-term live births, and use scRNA-seq to characterise the cell types present and their gene expression profiles. Additionally, we highlight the ability of this approach to detect significant gene expression changes across the second trimester of pregnancy in specific cell types, as well as identify changes in the proportions of cell types detected in AF across development. Overall, these findings are a proof-of-concept, highlighting the potential of this approach to deepen insight into both normal and abnormal human fetal development across the second trimester and, likely, later pregnancy.

## Methods

### Sample collection and preparation

AF samples were collected via clinically indicated amniocentesis performed on live ongoing pregnancies at a tertiary maternity hospital in Melbourne, VIC, Australia. AF was aspirated via a 22-gauge needle under ultrasound guidance. Five mL of fresh AF from each participant was retained for research purposes with written informed consent and transferred immediately to the research laboratory for processing. The AF was centrifuged at 300 × *g* for 10 minutes at 4°C, and the supernatant removed. The cell pellet was gently resuspended in 5 mL of Dulbecco’s phosphate-buffered saline (DPBS) and centrifuged at 300 × *g* for 10 minutes at 4°C, and the supernatant removed. For samples AF_1 and AF_2, the cell pellet was resuspended in DPBS at 1000 cells/µL and stored on ice until they were processed for sequencing. For the other samples, the cell pellet was cryopreserved in 90% fetal bovine serum (FBS) and 10% dimethyl sulfoxide (DMSO) until further use. This study was approved by the Human Research Ethics Committee of Mercy Health (2020-013).

### Single-cell RNA-sequencing

The cDNA preparation and library construction (end repair, A-tailing, adaptor ligation) was completed according to the Single Cell 3’ Protocol recommended by the manufacturer, 10x Genomics. The final libraries contain the P5 and P7 primers used in Illumina bridge amplification. Approximately 10,000 cells were loaded for cell capture using 10x Genomics Chromium machine. Single-cell capture was performed on fresh AF for samples AF_1 and AF_2, whereas the other samples (AF_3 to AF_7) were viably frozen prior to processing. Two captures were performed for sample AF_1 to recover a larger number of cells. Sequencing was carried out on the Illumina NextSeq 500 platform using the P2 100 cycle kit (400 million reads). Batch 1 comprised samples AF_1 and AF_2, batch 2 AF_3, batch 3 AF_4 and AF_5 and batch 4 AF_6 and AF_7.

### Dataset preprocessing

Full details of the bioinformatic analyses performed in this paper can be found at: https://oshlacklab.com/amniotic-fluid-single-cell-paper/

Samples AF_1 to AF_5 were processed using *Cell Ranger* 6.0.2 using human transcriptome reference version GRCh38 3.0.0 provided by *Cell Ranger*. All samples were also processed with *Cell Ranger*7.2.0 using the optimised human transcriptome reference described by Pool et al. (2023) [10]. R 4.5.0 was used to perform analysis using the *DropletUtils* 1.30.0 and *SingleCellExperiment* 1.32.0 R packages. For each capture, empty droplets were removed using the *emptyDrops()* function from the *DropletUtils* [11,12]. Analyses were restricted to cells retained in both pre-processing pipelines, enabling direct comparison of cell type annotation confidence at the single-cell level. To assess whether the optimised reference improved data quality, samples AF_1 to AF_5 were processed with both the standard *Cell Ranger* and optimised reference transcriptomes. The number of cells per sample, library size and number of detected genes per cell were compared before and after quality control filtering, using the *scater* 1.38.0 R package. During filtering, cells with a library size below 500, fewer than 300 detected genes or an outlying mitochondrial read proportion were removed. Finally, cells from both versions were annotated using *CellTypist* with the *Pan_Fetal_Human* model, and annotation confidence scores were compared.

For all subsequent analyses, the samples processed using the optimised reference were used as this was found to improve dataset quality. 11,454 cells remained across all samples after the quality filtering described above. Samples were classified into three groups by gestational age:

(1) early second trimester ≤ 19 weeks (n = 3: AF_3 [15 weeks], AF_5 [16 weeks], AF_7 [16 weeks])
(2) mid second trimester > 19 weeks and < 23 weeks (n = 1: AF_4 [20 weeks])
(3) late second trimester ≥ 23 weeks (n = 3: AF_1 [23 weeks], AF_6 [24 weeks], AF_2 [25 weeks])

### Cell type annotation and clustering

Cell type annotation was performed using the *CellTypist* 1.7.1 command line tool [13,14]. This was done with two different reference models: *Pan_Fetal_Human* and *Developing_Human_Organs* [15,16]. The *Pan_Fetal_Human* model places a greater emphasis on stromal and immune cell types [16] and the *Developing_Human_Organs* model places a greater emphasis on cell types from the lung, liver, stomach and intestine [15]. *CellTypist* applies a logistic regression model to classify cell types in a dataset using a reference model. Following cell type assignment, *CellTypist* implements an over-clustering approach with majority voting, ensuring that all cells within a subcluster share the same annotation (see Xu et al. (2023) and Domínguez Conde et al. (2022) for details) [13,14]. For this project, cell annotations obtained from both the *Pan_Fetal_Human* and *Developing_Human_Organs* reference models were examined and cells annotated by *CellTypist* as poor quality were removed, resulting in a total of 11,414 cells [15,16]. This is the number of cells used in all subsequent analyses on this dataset.

Reference-based cell annotation was followed by *SCTransform()* normalisation, anchor-based batch integration with *IntegrateData()* (using the first 30 dimensions, reciprocal PCA reduction and *k.anchor* = 20), dimensionality reduction and unsupervised clustering using the *Seurat* 4.4.0 R package [17–19]. The *clustree* 0.5.1 R package was used to visualise varying clustering resolutions [20]. To capture rare cell populations without over-clustering, the clustering approach consisted of three steps: low-resolution clustering, followed by subclustering of large clusters, and finally cluster refinement. For the low-resolution clusters, a resolution of 0.02 was used, producing seven broad clusters. The four clusters containing >500 cells were then subclustered at 0.3 resolution using *FindSubCluster().* Given the inherent similarity of T cell subtypes, the low-resolution T cell cluster was subclustered at 0.6 resolution by subsetting the dataset to these cells and then recomputing PCA, highly variable genes (HVGs) and batch integration steps. Within the T cell subclusters, distinct populations were identified by expression of canonical marker genes and using the top 20 marker genes of each subcluster (identified using *limma* and *treat*, as described below). T cell subclusters without distinct canonical markers were grouped into a single broader T cell cluster. Across the whole dataset, subclustering produced 32 clusters in total. Clusters annotated as the same cell type by both the *Developing_Human_Organs* and *Pan_Fetal_Human* references were then merged, giving 17 refined clusters that were used for all subsequent analyses.

Marker gene and Gene Ontology (GO) analysis was performed on each cluster to confirm its cell type identity. For marker gene analysis, counts were log₂-transformed with a pseudocount of 0.5 scaled to library size. Using the *limma* 3.66.0 R package, a design matrix accounting for the sample of origin was fitted with moderated *t*-tests incorporating mean–variance trend modelling and robust empirical Bayes shrinkage. Significant marker genes were identified with the *treat()* method, applying a log₂ fold-change threshold of 0.5 and Benjamini-Hochberg false discovery rate (FDR) < 0.05 [21–23]. The top 50 marker genes per cluster (ranked by t-statistic and filtered by log-fold change > 0) were then used to perform GO over-representation analysis using the *goana()* function from the *limma* R package [21]. Only terms from the Biological Process (BP) ontology and containing 200 or fewer genes were considered. Expression of canonical cell type marker genes, the top 20 marker genes for each cluster and cell type annotations provided by *CellTypist* were together used to label each of the 17 refined clusters.

### Cluster proportion analysis

The *sccomp* 2.2.0 R package was used to test for differences in cluster proportions between early and late gestational age groups [24]. The *sccomp_estimate()* function with the *bimodal_mean_variability_association* parameter set to true was used to fit the model. Outlier removal was performed using *sccomp_remove_outliers()* before performing hypothesis testing with *sccomp_test().* FDR < 0.05 was used to determine significant results.

### Trajectory analysis

Trajectory analysis was performed on clusters containing lung epithelium cells using the *Monocle3* 1.4.27 R package [25–27]. The low confidence epithelium cluster was used as the root for the pseudotime ordering. The resultant pseudotime values were mapped onto cells in the lung cluster and visualised using a UMAP. Trajectory analysis was also performed with the unknown cluster cells included to determine whether they are developmentally related. A trajectory UMAP for this was generated using the *plot_cells()* function from *Monocle3*.

### Differential gene expression in fetal skin cells

To examine differences in gene expression in fetal skin cells (Keratinocyte cluster) between the early and late gestational age groups, differential gene expression (DGE) was performed using a pseudobulk approach with the *RUVSeq* 1.44.0 and *edgeR* 4.8.2 R packages [28–30]. Sample AF_4 from the mid gestational age group was excluded from this analysis. Genes that could not be mapped to an Entrez ID, had zero counts or had a median log-CPM of ≤ 0 across the pseudobulked samples were removed prior to testing. The *RUVg()* function was used with housekeeping genes as negative control genes to identify factors of unwanted variation and the *k* parameter set to 1. The housekeeping genes used were the human *bulkRNAseqHK* list provided in the *scMerge* 1.26.0 R package [31,32]. DGE was performed using *edgeR* generalised linear model likelihood ratio test, taking into account fetal sex, gestational age group and the *W* factor estimated by *RUVSeq*. Genes with FDR < 0.05 were considered statistically significant.

### Gene set enrichment analysis

The *cameraPR()* function from *limma* [21,33] was used to perform Gene Set Enrichment Analysis (GSEA) on the Broad MSigDB v7.1 C2 Reactome and C5 gene sets [34–36] to identify enriched GO terms and Reactome pathways. The ranking statistic used was signed square root of the likelihood ratio. Genes from the Epidermal Differentiation Complex (EDC) were also tested for enrichment using *cameraPR()*. The EDC gene list was obtained from the ‘Gene group: Epidermal differentiation complex’ resource by HGNC [37,38].

### Comparison with an independent AF dataset

An AF scRNA-seq dataset generated by Gerli et al. (2024) was used for comparison (supplied pre-processed and integrated; obtained from NCBI Gene Expression Omnibus (GSE220994)) [1]. This dataset comprises 12 samples ranging from 15 to 34 weeks gestational age, with 34,568 cells in total. Of the 12 pregnancies (including 5 twin and 7 singleton gestations), 10 had a documented pathology, including twin-to-twin transfusion syndrome (n = 5), pathogenic chromosomal abnormalities (n = 2), and fetal structural abnormalities (n = 3). Two pregnancies were affected by more than one abnormality, and two involved fetal death. Two pregnancies had no recorded indication for amniocentesis. Most samples (n = 10) were obtained at the time of therapeutic amniodrainage (Figure 1A) [1]. We performed cell type annotation using the approach previously described for our dataset and cells annotated as poor quality or doublets were removed, resulting in 33,634 remaining cells. Clustering was also performed as described for our dataset, with 0.04 resolution used for broad clusters and 0.3 resolution for subclustering. After subclustering, 37 clusters were refined to 21 clusters, which were used in all subsequent analyses.

**Figure 1.**
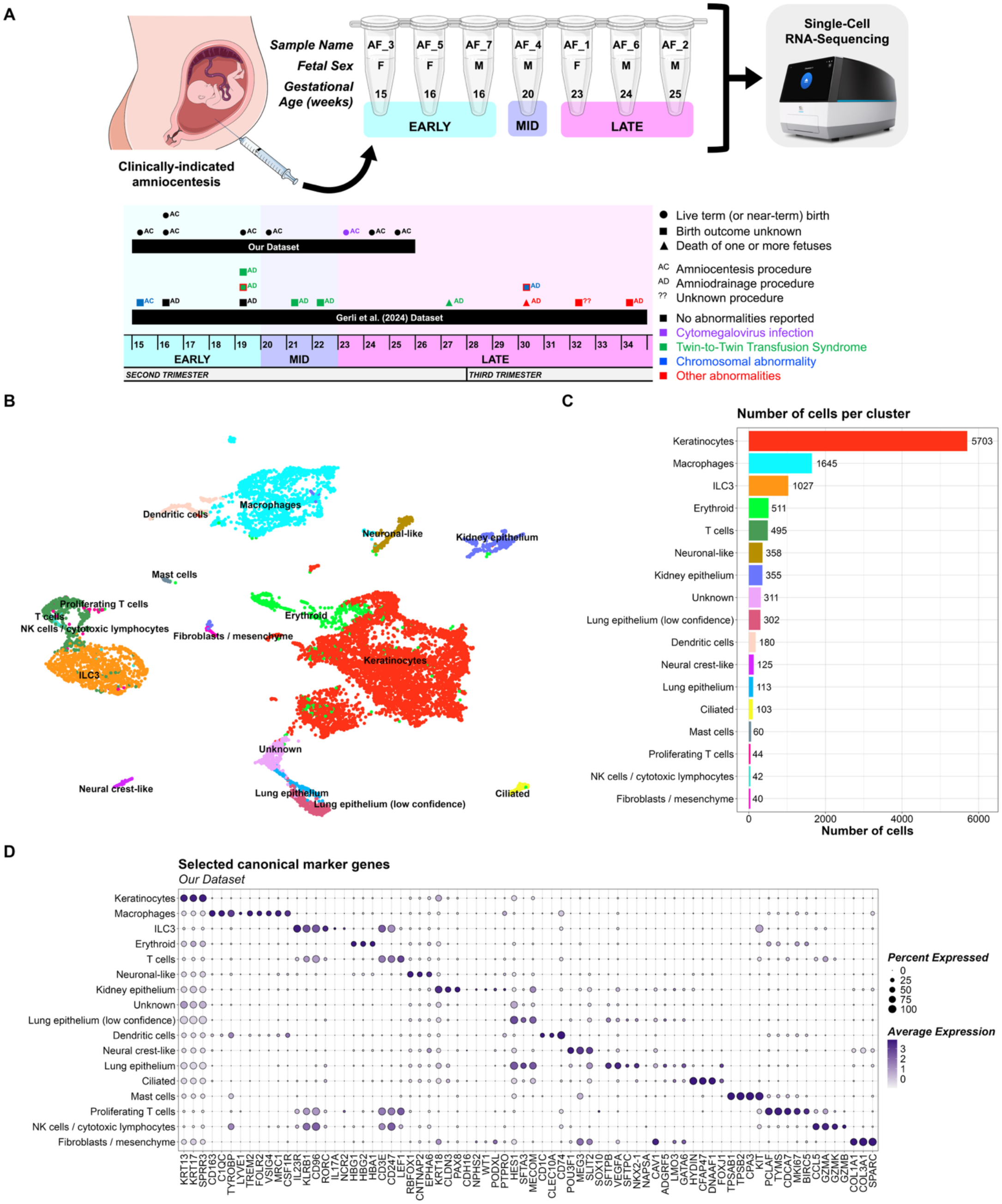
**(A)** Diagram overview of the samples in our dataset, including sample ID, gestational age and fetal sex, and how the gestational age, fetal abnormalities and birth outcomes of samples in our dataset and the Gerli et al. (2024) dataset compare [1]. Gestational age group (early = cyan, mid = lavender, late = magenta) for each sample is also shown. Point shape represents birth outcome, with circles for live term (or near-term) births, triangles for fetal death, and squares for samples without explicit reporting of birth outcome. Text next to each point describes the procedure performed during sample collection, with amniocentesis being performed for all samples in our dataset and most samples from the Gerli et al. (2024) dataset arising from amniodrainage procedures. Point colour represents any fetal abnormalities present in each sample, with black representing no abnormalities, purple representing cytomegalovirus infection, green representing twin-to-twin transfusion syndrome, blue representing chromosomal abnormalities, and red representing other abnormalities (congenital heart disease, oesophageal atresia, hydrops fetalis). Points with two colours represent those affected by multiple abnormalities. Graphics used in this figure were adapted from Servier Medical Art (https://smart.servier.com), licensed under CC BY 4.0 (https://creativecommons.org/licenses/by/4.0/). **(B)** UMAP plot of our dataset coloured by cluster label. **(C)** Number of cells per cluster in our dataset. **(D)** Dot plot of the expression of selected canonical cell type marker genes for each cluster in our dataset.

Samples in the Gerli et al. (2024) dataset were classified into three groups by gestational age that match the groups used for our dataset (early second trimester: n = 5; mid second trimester: n = 2; late second and third trimester: n = 5). As described for our dataset above, differences in the cluster proportions between early and late gestational age groups were compared using *sccomp*. DGE analysis in fetal skin cells (keratinocyte cluster) was also performed between the early and late gestational age groups as previously described, with k = 2 used for *RUVg*(). GSEA was then performed using the DGE results as described for our dataset.

To confirm whether the significant differentially expressed genes (DEGs) identified in our dataset were also changing across these developmental timepoints in the Gerli et al. (2024) dataset, the *cameraPR()* function from the *limma* R package was used to test whether the set of genes identified as significantly upregulated in the early gestational age group of our dataset (defined as genes with FDR < 0.05 and log fold-change > 0) were also highly ranked in the DEGs for the early group of the Gerli et al. (2024) dataset [21,33]. The same process was repeated for the significantly downregulated genes in the late gestational age group (defined as genes with FDR < 0.05 and log fold-change < 0) [21]. The results of this analysis were visualised using the *barcodeplot()* function from the *limma* R package.

Finally, the Gerli et al. (2024) dataset was integrated with ours to be provided together as a community resource. The *Seurat* 4.4.0 R package was used to perform normalisation via the *SCTransform()* method and dataset integration [17–19]. The integrated dataset contains 45,048 annotated cells and sample metadata.

## Results

### Single-cell profiling of amniotic fluid identified 11,414 cells

scRNA-seq was performed on AF samples collected via clinically indicated amniocentesis of seven pregnancies ranging from 15 to 25 weeks of gestation (Figure 1A). The indications for testing were increased risk of aneuploidy or fetal infection. All fetuses were structurally normal and euploid. One fetus (AF_1) was diagnosed with cytomegalovirus (CMV) infection but remained asymptomatic throughout pregnancy and at birth. All pregnancies resulted in live births of healthy term or near-term (≥36 weeks) infants (Supplementary Table 1).

AF is challenging to work with due to low cell retrieval and variable cell quality, thus we investigated whether the optimised human transcriptome reference developed by Pool et al. (2023) could increase cell numbers [10]. The optimised reference reduces counts sparsity by including counts from intronic reads, extended 3’ UTRs and genes overlapped by read-through transcripts [10]. Using this reference increased library sizes and number of detected genes per cell for each sample tested (Supplementary Figure 1A). More cells were also recovered per sample, with increases ranging from 11.9–382.3% before and 9.5–98.4% after quality filtering (Supplementary Figure 1A,B). Cell type annotations also improved in confidence compared to the standard *Cell Ranger* reference, as a greater proportion of cells per sample received higher cell type annotation probabilities (Supplementary Figure 1C). Consequently, the optimised reference was adopted for this study. After quality control, we identified a total of 11,454 cells across all samples. Cells annotated as poor-quality after the supervised cell type annotation described below were also removed, resulting in 11,414 cells for subsequent analyses. The number of cells per sample varied, with samples AF_7 and AF_5 having the most (3107) and fewest (250) cells, respectively (Supplementary Figure 2A). The samples in our dataset were of broadly similar quality and cell number to those described by Gerli et al. (2024), highlighting the challenging nature of performing scRNA-seq on AF (Supplementary Figure 2A).

### Amniotic fluid contains diverse fetal and immune cell populations

Supervised cell type annotation was performed using *CellTypist* with the *Pan_Fetal_Human* and *Developing_Human_Organs* reference models [13–16]. Unsupervised clustering identified 17 clusters that were broadly concordant with these annotations (Supplementary Figure 3E,F) and with marker gene and GO analyses (Supplementary Tables 2 and 3). Each cluster’s identity was then confirmed by expression of canonical marker genes (Figure 1B,D; Supplementary Figure 3). This revealed that AF is primarily composed of keratinocytes derived from fetal skin, which are annotated as keratinocytes or basal-like cells in the *Pan_Fetal_Human* and *Developing_Human_Organs* references, respectively (Figure 1B–D; Supplementary Figure 3). Various immune cell types were also identified, including macrophages, T cells, and lymphoid cells (Figure 1B–D; Supplementary Figure 3). These findings are consistent with previous literature on the cellular composition of AF [5,39]. Cells from the lung and kidney epithelium were also identified, suggesting a potential for using AF scRNA-seq to study the development and maturity of fetal organs. One cluster (Unknown) could not be confidently annotated, likely reflecting poor-quality cells and debris (Figure 1B–D). We also identified neuronal-like and neural crest-like clusters. The neuronal-like cluster was enriched for unexpected GO terms, including startle response and synaptic signalling (Supplementary Table 3), and its cells had low library sizes and few detected genes (Supplementary Figure 2B). Although neuroprogenitor cells may be present in AF, we cannot exclude that this cluster is driven by poor quality cells [8,40].

To test the reproducibility of these cell types, we applied the same analysis pipeline to an independent AF dataset, Gerli et al. (2024), comprising 12 pregnancies, 10 of which had fetal abnormalities. (Supplementary Figure 4) [1]. Despite these differences and the inclusion of later gestational ages (up to 34 weeks), the two datasets had broadly similar cell type compositions and integrated well (Figure 1B,C; Supplementary Figure 2C-E,3,4) [1]. Marker gene and GO analysis also corroborated the cell type annotations (Supplementary Table 4,5). In contrast with our data, the Gerli et al. (2024) dataset contained many additional immune cell types such as neutrophils, classical monocytes and other granulocytes (Supplementary Figure 4B). Furthermore, the Gerli et al. (2024) dataset also lacked the neuronal-like cluster observed in our dataset, further supporting that it may be an artefact of poor-quality cells (Supplementary Figure 4B). These discrepancies may reflect differences in sample preparation (e.g. cell sorting), fetal health and gestational age. Nevertheless, the core cell types were reproducible across both datasets, supporting AF as a source of diverse live fetal cells for single-cell study.

### Cell type proportions vary across gestational age

As AF contains cells shed from developing fetal tissues, we reasoned that its cellular composition might change as organs mature and thus compared cluster proportions between early (≤19 weeks) and late (≥23 weeks) gestational age groups. The ILC3 and lung epithelium clusters both showed significant changes in proportion between the early and late groups (FDR < 0.05; Figure 2A). ILC3 cells, which reside in the fetal skin and intestine, were abundant in the early samples but decreased in number and proportion in late samples (Figure 2A,B) [41,42]. Expression of canonical ILC3 marker genes did not differ notably between the early and late groups (Supplementary Figure 5C).

**Figure 2.**
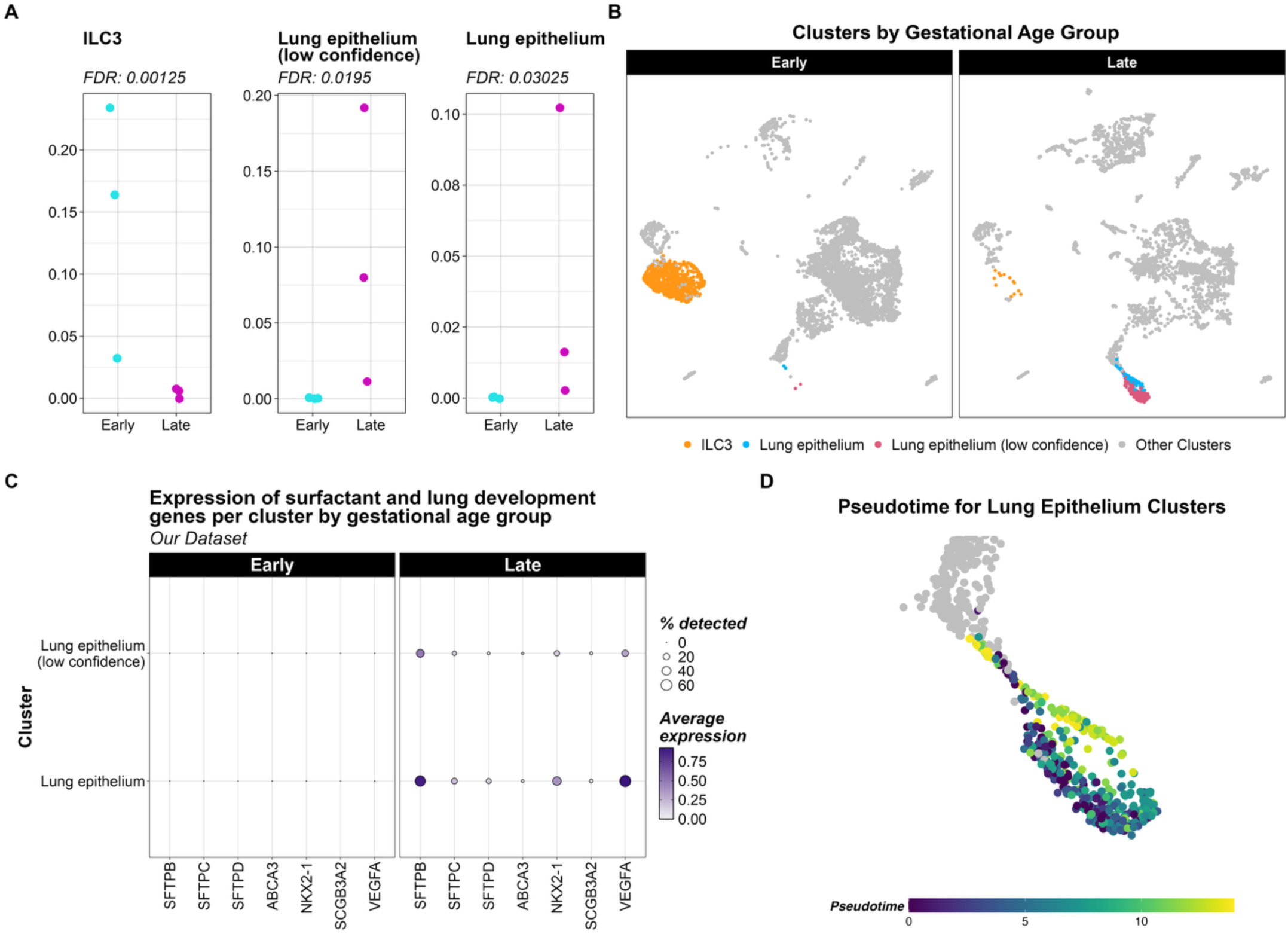
**(A)** Dot plots showing the cluster proportion values for each sample in the early and late gestational age groups for ILC3 and lung epithelium clusters in our dataset. The FDR result of the *sccomp* differential proportion test for each of these clusters is also shown. **(B)** UMAP plots showing the cells present in our dataset the early and late gestational age groups. ILC3 and lung epithelium clusters are coloured as these were found to have significant proportion differences between the early and late groups in *sccomp* testing. **(C)** Dot plot showing the expression of surfactant and lung development-related genes in lung epithelium clusters, split by gestational age group, for our dataset. Point colour reflects average expression value of that gene and point size reflects the percentage of cells it was detected in. **(D)** Zoomed in UMAP plot showing the lung epithelium clusters, coloured by pseudotime (as determined by *Monocle3*).

In contrast, the cell numbers and proportion of the lung epithelium clusters, increased with gestational age (Figure 2A,B). Based on the *Developing_Human_Organs* annotations, both lung epithelium clusters contained a mixture of mesenchyme subtype 4 and distal lung epithelium cells, with the higher confidence lung epithelium cluster containing a greater proportion of lung cells (Supplementary Figure 3F). In the Gerli et al. (2024) dataset, no clusters showed significant changes in proportion between the early and late groups, although the ILC3 and lung epithelium clusters followed trends similar to those in our dataset (Supplementary Figure 5B). Encouragingly, the cluster proportion trends in our dataset align with known fetal developmental processes. The greater proportion of lung epithelium cluster cells in the late group is consistent with lung development trends across this prenatal period (Figure 2A,B) [43]. Furthermore, we found higher expression of lung development-related and surfactant genes *SFTPB*, *SFTPC*, *SFTPD*, *VEGFA, ABCA3*, *SCGB3A2* and *NKX2-1* in the late gestational age group compared to the early group (Figure 2C). To determine whether the two lung epithelium clusters represent a developmental continuum, which could explain the “low confidence” label, we performed trajectory analysis. This showed that these clusters formed a trajectory, with the low confidence cluster rooted as the less differentiated state (Figure 2D). The Unknown cluster, examined to gain insight into its identity, remained separate from this trajectory (Supplementary Figure 5D).

Although the overall proportion of the T cell cluster did not change between age groups, its composition shifted (Supplementary Figure 5A,E,F). In the early group, this cluster mainly comprises type 1 innate T cells with fewer type 3 innate T cells (*Pan_Fetal_Human* reference). Conversely, in the late group, type 3 innate T cells comprise the majority of the cluster (Supplementary Figure 5F).

### Fetal skin cells show gestational age-related transcriptional maturation

To investigate how fetal skin matures across the second trimester, we compared gene expression in keratinocytes, the most abundant cell type in AF, between early and late gestational age groups using DGE analysis. In total, 1,304 significant DEGs were identified (FDR < 0.05), with 835 upregulated in the late gestational age group (Figure 3A). Keratin and keratin-associated genes *KRT4, KRT6C, KRT16, KRTDAP, KRT14, KRT31, KRT15, KRTAP19-7*, and *KRT24* were significantly upregulated in the late age group, and GSEA found that cornification- and keratinisation-related terms were over-represented, consistent with established knowledge of fetal skin development (Figure 3A,B) [44,45]. The EDC is located on chromosome 1q21 and contains genes involved in keratinocyte differentiation and establishment of the cornified envelope [46]. Similarly, EDC genes were enriched (p < 0.05) in the DEGs of the late group (Supplementary Figure 6A) [21,33].

**Figure 3.**
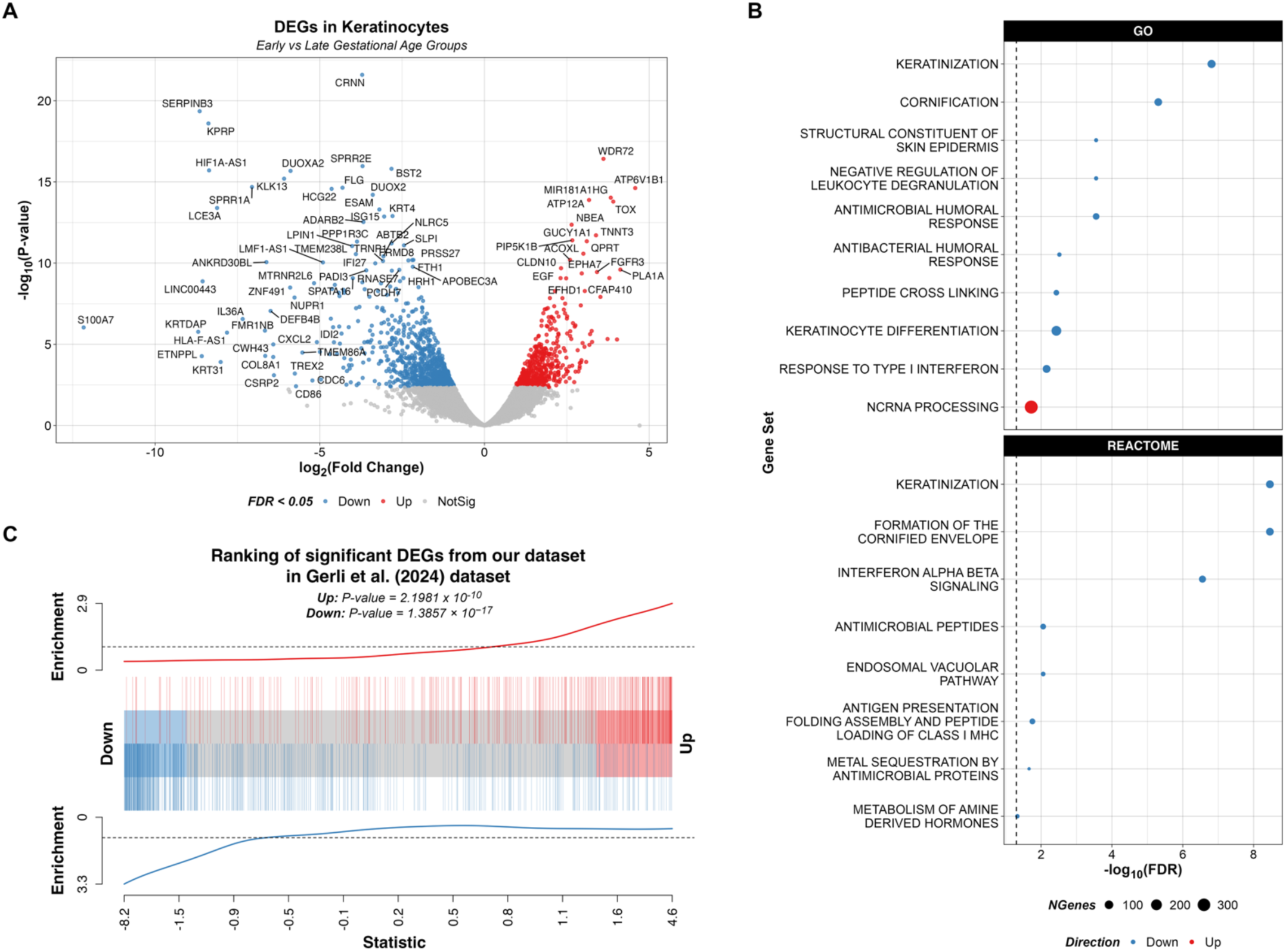
**(A)** Volcano plot showing the DEGs in fetal skin cells (keratinocytes) between the early and late gestational age groups in our dataset. Points are coloured by whether they were significantly upregulated in the early gestational age group (red; Up), significantly upregulated in the late gestational age group (blue; Down) or not significant (grey; NotSig). Points with an absolute log fold-change value above 5 or p-value below 0.000000001 were arbitrarily chosen to be labelled with their gene name. **(B)** Dot plots showing the GSEA enrichment results for DEGs identified for our dataset using the GO and Reactome gene sets. Points are coloured by whether they were significantly upregulated in the late gestational age group (blue; Down) or significantly upregulated in the early gestational age group (red, Up). Point size reflects number of genes in the tested gene set. **(C)** Barcode plot showing the enrichment of significant DEGs obtained by comparing fetal skin cells (keratinocytes) between early (red; Up) and late (blue; Down) gestational age groups in our dataset in the Gerli et al. (2024) dataset.

Applying the same DGE workflow to keratinocytes from the Gerli et al. (2024) dataset identified 22 and 157 significantly upregulated genes (FDR < 0.05) in the early and late age groups, respectively (Supplementary Figure 6B). GSEA highlighted changes in haptoglobin, haemoglobin, apoptosis, gas transport and receptor-related pathways that were not specific to skin development (Supplementary Figure 6C) and there was no significant enrichment of EDC genes (Supplementary Figure 6A). This likely reflects the greater clinical heterogeneity in their pathological cohort and the higher gestational age range in their late group. We therefore used competitive gene set testing to test whether the age-related changes identified in fetal skin cells from our healthy dataset were recapitulated in the Gerli et al. (2024) dataset. DEGs from our early and late groups were significantly enriched in the corresponding age groups of the Gerli et al. (2024) data (p < 0.05 for both), supporting the age-related changes identified in our dataset (Figure 3C) [21,33]. These findings establish the feasibility of DGE analysis in AF scRNA-seq data, with gestational age-related expression changes that reflect known skin development.

## Discussion

AF is a unique biofluid that provides access to live fetal cells from ongoing pregnancies, enabling the study of human development across timepoints inaccessible through terminated pregnancies. This work presents, to our knowledge, the first scRNA-seq atlas of AF from healthy second-trimester pregnancies, comprising seven samples from singleton pregnancies that all resulted in live births with no major congenital abnormalities (Supplementary Table 1). Our healthy reference set, together with our comparative analysis of the pathological cohort from Gerli et al. (2024) [1], offers proof-of-concept that AF scRNA-seq can capture gestational age-related changes in both cell type proportions and DEGs.

Keratinocytes were the dominant cell type in AF (Figure 1B,C), consistent with the skin forming the outer lining of the fetus in direct contact with AF and this was reproduced in the Gerli et al. (2024) dataset (Supplementary Figure 4A,B). DGE analysis revealed extensive changes in these cells across the second trimester, including increasing expression of various keratin and keratin-associated genes (Figure 3A). Some of these, such as *KRT4,* are expressed in mucosal and oesophageal epithelia, plausibly reflecting cells shed from these surfaces as the fetus swallows [47]. EDC genes were also significantly enriched in the late group (23 to 25 weeks; Supplementary Figure 6A). These are involved in keratinocyte differentiation and development of the cornified envelope, consistent with keratinisation commencing from 14 weeks and rapid skin maturation occurring around 24 weeks [44,48]. Although one late age group sample (AF_1) was CMV positive, CMV is not known to affect the skin and the infant was born asymptomatic, so it is unlikely to have affected the DGE results [49]. Together, these findings show that key developmental processes within a cellular compartment can be captured in AF.

In the Gerli et al. (2024) dataset, the keratinocyte DEGs detected in our healthy cohort were significantly enriched in the corresponding age groups (Figure 3C). However, performing DGE analysis on the Gerli et al. (2024) dataset on its own did not recover skin development-associated terms (Supplementary Figure 6C). We hypothesise that the heterogeneity of their largely abnormal pregnancies reduced power to detect developmental signals, which were recovered using a more targeted gene set test. Similarly, EDC genes were not significantly enriched in the Gerli et al. (2024) late age group, instead trending non-significantly towards the early group (Supplementary Figure 6A). The EDC drives terminal epithelial differentiation to form the stratum corneum, which begins forming around hair follicles at 18 weeks and extends across the interfollicular epidermis during weeks 23 to 26 [46,48]. There is also some evidence that the EDC is expressed in the embryonic periderm, which is absent after 23 weeks [46,48]. The Gerli et al. (2024) late group (27 to 34 weeks) falls outside the main window of stratum corneum development, whereas our late group (23 to 25 weeks) falls within it, which may contribute to the discrepancy (Figure 1A) [46,48].

We identified numerous immune cell types in AF, including macrophages, ILC3 cells and various T cell populations (Figure 1B,C), consistent with immune processes previously reported in the second-trimester AF cell-free transcriptome [7,8,50]. Similar immune populations were present in the Gerli et al. (2024) dataset, except for neutrophils, classical monocytes and other granulocytes, which were abundant in their data but absent from ours (Supplementary Figure 4A,B). This may reflect inflammatory conditions in these largely abnormal pregnancies, their later gestational ages, or differences in sample collection and processing. Indeed, as AF from normal pregnancies is a low-inflammatory environment [5], strong innate immune responses occur with infection such as CMV [7], and pro-inflammatory cytokines are overexpressed in spontaneous preterm birth [50]. A healthy reference such as ours is therefore essential for interpreting immune changes in disease. Within our dataset, the T cell cluster shifted from predominantly type 1 to type 3 innate T cells across gestation (Supplementary Figure 5F). Suo et al. (2022) described these as developmental-specific unconventional T cells but studied only early gestation [16], so their roles later in development warrant further study. Finally, *LYVE1^hi^* macrophages regulate morphogenesis of fetal skin and dermal vasculature [41,51]. Given that these macrophages (all annotated as *LYVE1^hi^* with the *Pan_Fetal_Human* reference; Supplementary Figure 3B) and keratinocytes were among the most abundant cell types in our dataset, AF may offer a means to study crosstalk between these two cell types.

The ILC3 and lung epithelium clusters changed significantly in proportion between early and late gestation (Figure 2A). The decrease in ILC3s is supported by evidence suggesting that ILCs are most abundant in the second trimester before decreasing [5,42,52]. Although sample AF_1 was CMV positive, its low ILC3 proportion was consistent with the other late group samples. Conversely, lung epithelium cells increased in the late group (Figure 2A), including a less differentiated, possibly transitional population as suggested by trajectory analysis (Figure 2D). The lung epithelium cells also had greater expression of surfactant and lung development-related genes in the late group (Figure 2C). Our early (15-16 weeks) and late groups (23-25 weeks) span the canalicular stage of fetal lung development (16-26 weeks) during which alveolar ducts and respiratory bronchioli develop, suggesting that AF scRNA-seq could be used to monitor fetal lung maturation. If confirmed in larger studies, this could have major clinical applications, such as predicting postnatal outcomes of extreme prematurity or assessing fetuses with congenital anomalies associated with pulmonary hypoplasia, such as congenital diaphragmatic hernia. Interpretation is limited by the small number of samples per group (n = 3) and unequal fetal sex balance (early = two female, one male; late = one female, two male). However, as lung maturation occurs more rapidly in female fetuses [53,54], the male-biased late group would, if anything, underestimate the increase in lung epithelium, so sex differences are unlikely to explain this result.

The *Pan_Fetal_Human* and *Developing_Human_Organs* references used for *CellTypist* annotation contain fetal tissue from 6 to 19 and 9 to 23 weeks gestational age, respectively [13–16], whereas our samples and the Gerli et al. (2024) samples span 15 to 25 and 15 to 34 weeks, respectively. Given the extent of fetal development across these timepoints, the gene expression profiles of many cell types in AF likely differ from these references. We therefore confirmed *CellTypist* labels with unsupervised clustering, marker gene analysis and GO analyses, which were generally concordant (Supplementary Figure 3,4; Supplementary Table 2-5).

For some clusters, however, unexpected GO terms emerged, such as neural development terms (Supplementary Table 3). These may be driven by poor-quality droplets, broader developmental pathway activity, or ambient RNA contamination. Ambient RNA is background RNA, typically from lysed cells, that is captured in droplets alongside intact cells. AF is a complex biofluid rich in debris, extracellular vesicles and cell-free RNA [7,8,50,55–57], including organ-specific transcripts from tissues such as the brain [55–57], so whether these GO terms reflect intact neuroprogenitor cells or ambient RNA remains unclear. Keratinocytes were the primary source of ambient contamination, with their marker genes detected across all other cell types (Figure 1D; Supplementary Figure 4C), consistent with abundant cell types with large library sizes dominating the ambient RNA profile (Supplementary Figure 2B) [58]. Such ambient RNA contamination is difficult to correct because the ambient profile resembles a genuine cell type, and there is little guidance on using ambient-corrected counts in between-conditions DGE analyses [58–64]. Given the risk of data distortion, the challenges of AF data, and our DGE focus on keratinocytes (the main source of ambient RNA), we deliberately chose not to apply ambient correction [11,60,61]. It should be noted that the Gerli et al. (2024) dataset was ambient-corrected using *CellBender* [65], which may contribute to differences between the datasets.

Sample quality is a major challenge for AF scRNA-seq, as cellular debris which accumulates with gestational age, causes variability in the number and quality of captured cells, potentially making later samples more challenging to process. Gerli et al. (2024) used FACS to select intact cells, yet our unsorted samples were of comparable quality (Supplementary Figure 2A). We also trialled nuclei extraction and FACS sorting of one fresh sample but found no clear benefit (data not shown), although other wet lab optimisations or using fixed-cell scRNA-seq protocols such as 10x Flex may improve capture of diverse cell types. Our cryopreserved AF samples were not of lower quality than the two fresh AF samples (Supplementary Figure 7A). This is consistent with reports that cryopreservation does not alter scRNA-seq transcriptional profiles [66], and cryopreservation offers clear logistical advantages over processing fresh AF. As both fresh samples were in the late group, processing method was partially confounded with gestational age. MDS plots of housekeeping gene expression separated fresh from frozen samples (Supplementary Figure 7B), indicating a processing effect. Our analysis approach using *RUVSeq* with housekeeping genes is designed to estimate and remove precisely this variation in our DGE analysis [28]. In contrast, cluster proportions separated primarily by gestational age group, with processing method contributing to a lesser extent (Supplementary Figure 7C).

Our work provides proof-of-concept findings, and replication in larger, more balanced cohorts is needed to further explore these biological observations. Beyond this, a healthy reference atlas opens opportunities to study prenatal disease. For example, AF_1 in our dataset was CMV-positive but showed no structural abnormalities on ultrasound and resulted in the term birth of a healthy, asymptomatic infant (Supplementary Table 1); examining AF from symptomatic CMV infections could provide unique insights into how infection alters fetal cell composition and gene expression. Similarly, AF scRNA-seq could be used to map the effects of chromosomal abnormalities across gestation, and copy number variant inference developed for tumour scRNA-seq, such as *inferCNV* [67], could be evaluated for detecting gene expression perturbations in aneuploid pregnancies.

Finally, given the DOHaD hypothesis [3], longitudinal studies that begin with AF scRNA-seq and follow individuals across their lifespan could link fetal cellular states to long-term health outcomes.

## Conclusion

AF scRNA-seq captures a diverse range of live fetal cell types and reveals gestational age-related changes in cell composition and gene expression, including signatures of lung maturation and skin development that were broadly reproduced in an independent cohort. This atlas provides a healthy reference for second-trimester development. It establishes AF scRNA-seq as a feasible approach for studying live fetal development and, in future, the cellular changes underlying prenatal disease.

## Code Availability

All code used to perform the analyses and generate the figures presented in this manuscript can be found at https://oshlacklab.com/amniotic-fluid-single-cell-paper/.

## Data Availability

Raw and processed single-cell count data for our dataset are available from Zenodo in *Cell Ranger* output and *Seurat* formats at https://doi.org/10.5281/zenodo.23655939. The integrated dataset combining our dataset and the Gerli et al. (2024) dataset [1] is also provided in *Seurat* format in this Zenodo record. Raw sequencing data (FASTQ) are not publicly available as per ethics approval.

## Author Contributions

### CRediT Author Contributions

- Conceptualization: LH, AO
- Funding acquisition: LH (CIA), NH, BP, AO, JM
- Resources: LH, NH, IA, SB, BP, SG
- Methodology: LH, SG, BP, JM, AO, CG
- Investigation (clinical – sample collection, processing, sequencing): IA, SB, SG, BP, NH
- Investigation (bioinformatics): CG, JM, AO, LN
- Writing – original draft: CG
- Writing – review & editing: All authors
- Supervision: LH, AO, JM

J.M., L.H. and A.O. are joint senior authors. L.H. provided senior clinical leadership, obtained funding as Chief Investigator A, oversaw sample acquisition, and provided overall study coordination, while J.M. and A.O. provided senior bioinformatics leadership and supervised the computational analyses.

## Acknowledgements

The authors thank the clinical and research staff at the Mercy Hospital for Women Department of Perinatal Medicine and the University of Melbourne Department of Obstetrics, Gynaecology and Newborn Health for infrastructure support and assistance with participant recruitment, sample collection and processing. We also thank the authors of Gerli et al. (2024) for providing additional information on their datasets to support our analysis.

## References

1. Gerli MFM, Calà G, Beesley MA, Sina B, Tullie L, Sun KY, et al. Single-cell guided prenatal derivation of primary fetal epithelial organoids from human amniotic and tracheal fluids. Nat Med. Nature Publishing Group; 2024;30:875–87. 10.1038/s41591-024-02807-z

2. Haniffa M, Taylor D, Linnarsson S, Aronow BJ, Bader GD, Barker RA, et al. A roadmap for the Human Developmental Cell Atlas. Nature. Nature Publishing Group; 2021;597:196–205. 10.1038/s41586-021-03620-1

3. Lacagnina S. The Developmental Origins of Health and Disease (DOHaD). Am J Lifestyle Med. 2019;14:47–50. 10.1177/1559827619879694

4. Gerrelli D, Lisgo S, Copp AJ, Lindsay S. Enabling research with human embryonic and fetal tissue resources. Dev Camb Engl. 2015;142:3073–6. 10.1242/dev.122820

5. Gomez-Lopez N, Romero R, Xu Y, Miller D, Leng Y, Panaitescu B, et al. The Immunophenotype of Amniotic Fluid Leukocytes in Normal and Complicated Pregnancies. Am J Reprod Immunol N Y N 1989. 2018;79:e12827. 10.1111/aji.12827

6. Zwemer LM, Bianchi DW. The Amniotic Fluid Transcriptome as a Guide to Understanding Fetal Disease. Cold Spring Harb Perspect Med. 2015;5:a023101. 10.1101/cshperspect.a023101

7. Atukorala I, Hannan N, Hui L. Immersed in a reservoir of potential: amniotic fluid-derived extracellular vesicles. J Transl Med. 2024;22:348. 10.1186/s12967-024-05154-2

8. Hui L, Slonim DK, Wick HC, Johnson KL, Bianchi DW. The Amniotic Fluid Transcriptome: A Source of Novel Information About Human Development. Obstet Gynecol. 2012;119:111–8. 10.1097/AOG.0b013e31823d4150

9. Ghi T, Sotiriadis A, Calda P, Da Silva Costa F, Raine-Fenning N, Alfirevic Z, et al. ISUOG Practice Guidelines: invasive procedures for prenatal diagnosis. Ultrasound Obstet Gynecol. 2016;48:256–68. 10.1002/uog.15945

10. Pool A-H, Poldsam H, Chen S, Thomson M, Oka Y. Recovery of missing single-cell RNA-sequencing data with optimized transcriptomic references. Nat Methods. Nature Publishing Group; 2023;20:1506–15. 10.1038/s41592-023-02003-w

11. Lun ATL, Riesenfeld S, Andrews T, Dao TP, Gomes T, Marioni JC, et al. EmptyDrops: distinguishing cells from empty droplets in droplet-based single-cell RNA sequencing data. Genome Biol. 2019;20:63. 10.1186/s13059-019-1662-y

12. Griffiths JA, Richard AC, Bach K, Lun ATL, Marioni JC. Detection and removal of barcode swapping in single-cell RNA-seq data. Nat Commun. Nature Publishing Group; 2018;9:2667. 10.1038/s41467-018-05083-x

13. Xu C, Prete M, Webb S, Jardine L, Stewart BJ, Hoo R, et al. Automatic cell-type harmonization and integration across Human Cell Atlas datasets. Cell. Elsevier; 2023;186:5876–5891.e20. 10.1016/j.cell.2023.11.026

14. Domínguez Conde C, Xu C, Jarvis LB, Rainbow DB, Wells SB, Gomes T, et al. Cross-tissue immune cell analysis reveals tissue-specific features in humans. Science. American Association for the Advancement of Science; 2022;376:eabl5197. 10.1126/science.abl5197

15. Yu Q, Kilik U, Holloway EM, Tsai Y-H, Harmel C, Wu A, et al. Charting human development using a multi-endodermal organ atlas and organoid models. Cell. Elsevier; 2021;184:3281–3298.e22. 10.1016/j.cell.2021.04.028

16. Suo C, Dann E, Goh I, Jardine L, Kleshchevnikov V, Park J-E, et al. Mapping the developing human immune system across organs. Science. American Association for the Advancement of Science; 2022;376:eabo0510. 10.1126/science.abo0510

17. Choudhary S, Satija R. Comparison and evaluation of statistical error models for scRNA-seq. Genome Biol. 2022;23:27. 10.1186/s13059-021-02584-9

18. Hafemeister C, Satija R. Normalization and variance stabilization of single-cell RNA-seq data using regularized negative binomial regression. Genome Biol. 2019;20:296. 10.1186/s13059-019-1874-1

19. Hao Y, Hao S, Andersen-Nissen E, Mauck WM, Zheng S, Butler A, et al. Integrated analysis of multimodal single-cell data. Cell. 2021;184:3573–3587.e29. 10.1016/j.cell.2021.04.048

20. Zappia L, Oshlack A. Clustering trees: a visualization for evaluating clusterings at multiple resolutions. GigaScience. 2018;7:giy083. 10.1093/gigascience/giy083

21. Ritchie ME, Phipson B, Wu D, Hu Y, Law CW, Shi W, et al. limma powers differential expression analyses for RNA-sequencing and microarray studies. Nucleic Acids Res. 2015;43:e47. 10.1093/nar/gkv007

22. McCarthy DJ, Smyth GK. Testing significance relative to a fold-change threshold is a TREAT. Bioinformatics. 2009;25:765–71. 10.1093/bioinformatics/btp053

23. Benjamini Y, Hochberg Y. Controlling the False Discovery Rate: A Practical and Powerful Approach to Multiple Testing. J R Stat Soc Ser B Methodol. [Royal Statistical Society, Oxford University Press]; 1995;57:289–300.

24. Mangiola S, Roth-Schulze AJ, Trussart M, Zozaya-Valdés E, Ma M, Gao Z, et al. sccomp: Robust differential composition and variability analysis for single-cell data. Proc Natl Acad Sci U S A. 2023;120:e2203828120. 10.1073/pnas.2203828120

25. Cao J, Spielmann M, Qiu X, Huang X, Ibrahim DM, Hill AJ, et al. The single cell transcriptional landscape of mammalian organogenesis. Nature. 2019;566:496–502. 10.1038/s41586-019-0969-x

26. Qiu X, Mao Q, Tang Y, Wang L, Chawla R, Pliner HA, et al. Reversed graph embedding resolves complex single-cell trajectories. Nat Methods. 2017;14:979–82. 10.1038/nmeth.4402

27. Trapnell C, Cacchiarelli D, Grimsby J, Pokharel P, Li S, Morse M, et al. Pseudo-temporal ordering of individual cells reveals dynamics and regulators of cell fate decisions. Nat Biotechnol. 2014;32:381–6. 10.1038/nbt.2859

28. Risso D, Ngai J, Speed TP, Dudoit S. Normalization of RNA-seq data using factor analysis of control genes or samples. Nat Biotechnol. Nature Publishing Group; 2014;32:896–902. 10.1038/nbt.2931

29. Robinson MD, McCarthy DJ, Smyth GK. edgeR: a Bioconductor package for differential expression analysis of digital gene expression data. Bioinformatics. 2010;26:139–40. 10.1093/bioinformatics/btp616

30. McCarthy DJ, Chen Y, Smyth GK. Differential expression analysis of multifactor RNA-Seq experiments with respect to biological variation. Nucleic Acids Res. 2012;40:4288–97. 10.1093/nar/gks042

31. Lin Y, Ghazanfar S, Wang KYX, Gagnon-Bartsch JA, Lo KK, Su X, et al. scMerge leverages factor analysis, stable expression, and pseudoreplication to merge multiple single-cell RNA-seq datasets. Proc Natl Acad Sci. Proceedings of the National Academy of Sciences; 2019;116:9775–84. 10.1073/pnas.1820006116

32. Lin Y, Ghazanfar S, Strbenac D, Wang A, Patrick E, Lin DM, et al. Evaluating stably expressed genes in single cells. GigaScience. 2019;8:giz106. 10.1093/gigascience/giz106

33. Wu D, Smyth GK. Camera: a competitive gene set test accounting for inter-gene correlation. Nucleic Acids Res. 2012;40:e133. 10.1093/nar/gks461

34. Liberzon A, Birger C, Thorvaldsdóttir H, Ghandi M, Mesirov JP, Tamayo P. The Molecular Signatures Database (MSigDB) hallmark gene set collection. Cell Syst. 2015;1:417–25. 10.1016/j.cels.2015.12.004

35. Liberzon A, Subramanian A, Pinchback R, Thorvaldsdóttir H, Tamayo P, Mesirov JP. Molecular signatures database (MSigDB) 3.0. Bioinformatics. 2011;27:1739–40. 10.1093/bioinformatics/btr260

36. Subramanian A, Tamayo P, Mootha VK, Mukherjee S, Ebert BL, Gillette MA, et al. Gene set enrichment analysis: A knowledge-based approach for interpreting genome-wide expression profiles. Proc Natl Acad Sci. Proceedings of the National Academy of Sciences; 2005;102:15545–50. 10.1073/pnas.0506580102

37. Seal RL, Braschi B, Gray K, Jones TEM, Tweedie S, Haim-Vilmovsky L, et al. Genenames.org: the HGNC resources in 2023. Nucleic Acids Res. 2023;51:D1003–9. 10.1093/nar/gkac888

38. Epidermal differentiation complex | HUGO Gene Nomenclature Committee [Internet]. [cited 2025 Aug 27]. https://www.genenames.org/data/genegroup/#!/group/1354. Accessed 27 Aug 2025

39. Hoehn H, Salk D. Chapter 2. Morphological and Biochemical Heterogeneity of Amniotic Fluid Cells in Culture. In: Latt SA, Darlington GJ, editors. Methods Cell Biol [Internet]. Academic Press; 1982. p. 11–34. 10.1016/S0091-679X(08)61362-X

40. Bossolasco P, Montemurro T, Cova L, Zangrossi S, Calzarossa C, Buiatiotis S, et al. Molecular and phenotypic characterization of human amniotic fluid cells and their differentiation potential. Cell Res. Nature Publishing Group; 2006;16:329–36. 10.1038/sj.cr.7310043

41. Gopee NH, Winheim E, Olabi B, Admane C, Foster AR, Huang N, et al. A prenatal skin atlas reveals immune regulation of human skin morphogenesis. Nature. Nature Publishing Group; 2024;635:679–89. 10.1038/s41586-024-08002-x

42. Marquardt N, Ivarsson MA, Sundström E, Åkesson E, Martini E, Eidsmo L, et al. Fetal CD103+ IL-17–Producing Group 3 Innate Lymphoid Cells Represent the Dominant Lymphocyte Subset in Human Amniotic Fluid. J Immunol. 2016;197:3069–75. 10.4049/jimmunol.1502204

43. Rehman S, Bacha D. Embryology, Pulmonary. StatPearls [Internet]. Treasure Island (FL): StatPearls Publishing; 2025 [cited 2025 Apr 10]. http://www.ncbi.nlm.nih.gov/books/NBK544372/. Accessed 10 Apr 2025

44. Larson BJ, Longaker MT, Lorenz HP. Scarless Fetal Wound Healing: A Basic Science Review. Plast Reconstr Surg. 2010;126:1172–80. 10.1097/PRS.0b013e3181eae781

45. Coolen NA, Schouten KCWM, Middelkoop E, Ulrich MMW. Comparison between human fetal and adult skin. Arch Dermatol Res. 2010;302:47–55. 10.1007/s00403-009-0989-8

46. Holthaus KB, Eckhart L. Development-Associated Genes of the Epidermal Differentiation Complex (EDC). J Dev Biol. 2024;12:4. 10.3390/jdb12010004

47. KRT4 keratin 4 [Homo sapiens (human)] - Gene - NCBI [Internet]. [cited 2025 Aug 27]. https://www.ncbi.nlm.nih.gov/gene?Db=gene&Cmd=DetailsSearch&Term=3851. Accessed 27 Aug 2025

48. Visscher M, Narendran V. The Ontogeny of Skin. Adv Wound Care. 2014;3:291–303. 10.1089/wound.2013.0467

49. Pass RF, Arav-Boger R. Maternal and fetal cytomegalovirus infection: diagnosis, management, and prevention. F1000Research. 2018;7:255. 10.12688/f1000research.12517.1

50. Bhatti G, Romero R, Gomez-Lopez N, Pique-Regi R, Pacora P, Jung E, et al. The amniotic fluid cell-free transcriptome in spontaneous preterm labor. Sci Rep. Nature Publishing Group; 2021;11:13481. 10.1038/s41598-021-92439-x

51. Blériot C, Chakarov S, Ginhoux F. Determinants of Resident Tissue Macrophage Identity and Function. Immunity. Elsevier; 2020;52:957–70. 10.1016/j.immuni.2020.05.014

52. Mendes J, Areia AL, Rodrigues-Santos P, Santos-Rosa M, Mota-Pinto A. Innate Lymphoid Cells in Human Pregnancy. Front Immunol [Internet]. 2020;Volume 11-2020. https://www.frontiersin.org/journals/immunology/articles/10.3389/fimmu.2020.551707

53. Torday JS, Nielsen HC, Fencl MDM, Avery ME. Sex Differences in Fetal Lung Maturation. Am Rev Respir Dis. 1981;123:205–8. 10.1164/arrd.1981.123.2.205

54. Tondo P, Meschi C, Mantero M, Scioscia G, Siciliano M, Bradicich M, et al. Sex and gender differences during the lung lifespan: unveiling a pivotal impact. Eur Respir Rev [Internet]. European Respiratory Society; 2025 [cited 2026 Sept 28];34. 10.1183/16000617.0121-2024

55. Hui L, Tong S, Kaitu’u-Lino TJ, Hannan NJ. A comparison of sample collection methods for quantifying cell-free fetal neurodevelopment transcripts in amniotic fluid. BMC Res Notes. 2016;9:335. 10.1186/s13104-016-2146-8

56. Hui L, De Catte L, Beard S, Maksimovic J, Vora NL, Oshlack A, et al. RNA-Seq of amniotic fluid cell-free RNA: a discovery phase study of the pathophysiology of congenital cytomegalovirus infection. Am J Obstet Gynecol. 2022;227:634.e1–634.e12. 10.1016/j.ajog.2022.05.035

57. Park HJ, Cho HY, Cha DH. The Amniotic Fluid Cell-Free Transcriptome Provides Novel Information about Fetal Development and Placental Cellular Dynamics. Int J Mol Sci. 2021;22:2612. 10.3390/ijms22052612

58. Caglayan E, Liu Y, Konopka G. Neuronal ambient RNA contamination causes misinterpreted and masked cell types in brain single-nuclei datasets. Neuron. Elsevier; 2022;110:4043–4056.e5. 10.1016/j.neuron.2022.09.010

59. Schaefer NK, Pavlovic BJ, Schmitz MT, Pollen AA. CellBouncer, a unified toolkit for single-cell demultiplexing and ambient RNA analysis, reveals hominid mitochondrial incompatibilities. Cell Genomics [Internet]. Elsevier; 2026 [cited 2026 Oct 21];6. 10.1016/j.xgen.2026.101275

60. Cargnelli CB, Nielsen JV, Madsen JGS. Benchmarking computational decontamination of ambient RNA. Front Bioinforma [Internet]. Frontiers; 2026 [cited 2026 Oct 21];6. 10.3389/fbinf.2026.1844838

61. Schroeder L, Gerber S, Ruffini N. Benchmarking ambient RNA removal across droplet and well-plate platforms reveals artificial count generation as a critical failure mode of scAR and CellClear [Internet]. bioRxiv; 2026 [cited 2026 May 18]. p. 2026.04.08.717130. 10.64898/2026.04.08.717130

62. Janssen P, Kliesmete Z, Vieth B, Adiconis X, Simmons S, Marshall J, et al. The effect of background noise and its removal on the analysis of single-cell expression data. Genome Biol. 2023;24:140. 10.1186/s13059-023-02978-x

63. Nie Y, Zhang C, Tan J, Chen X, Yuan W, Luo X, et al. Systematic benchmarking of ambient RNA decontamination tools to advance precision in single-cell transcriptomic analysis. Nat Commun. Nature Publishing Group; 2026;17:10604. 10.1038/s41467-026-77458-4

64. Berg M, Petoukhov I, van den Ende I, Meyer KB, Guryev V, Vonk JM, et al. FastCAR: fast correction for ambient RNA to facilitate differential gene expression analysis in single-cell RNA-sequencing datasets. BMC Genomics. 2023;24:722. 10.1186/s12864-023-09822-3

65. Fleming SJ, Chaffin MD, Arduini A, Akkad A-D, Banks E, Marioni JC, et al. Unsupervised removal of systematic background noise from droplet-based single-cell experiments using CellBender. Nat Methods. Nature Publishing Group; 2023;20:1323–35. 10.1038/s41592-023-01943-7

66. Guillaumet-Adkins A, Rodríguez-Esteban G, Mereu E, Mendez-Lago M, Jaitin DA, Villanueva A, et al. Single-cell transcriptome conservation in cryopreserved cells and tissues. Genome Biol. 2017;18:45. 10.1186/s13059-017-1171-9

67. Tickle T, Tirosh I, Georgescu C, Brown M, Haas B. inferCNV of the Trinity CTAT Project. [Internet]. Klarman Cell Observatory, Broad Institute of MIT and Harvard, Cambridge, MA, USA.; 2019. https://github.com/broadinstitute/inferCNV

