## Supplementary Figures for "Single-cell RNA-seq of amniotic fluid provides unique insight into fetal development"

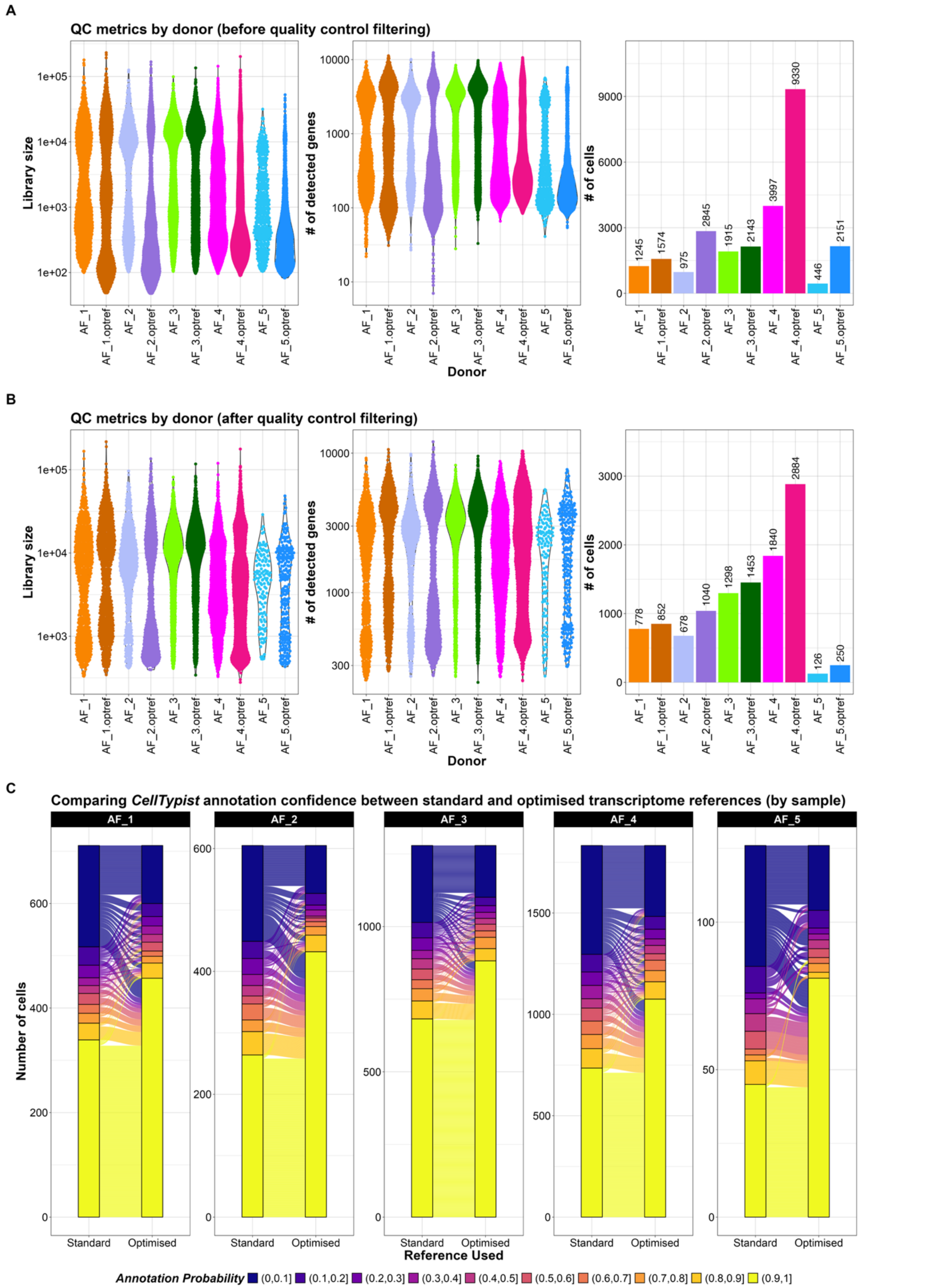

**Supplementary Figure 1. (A)** Comparison of quality control metrics after empty droplet removal of samples processed using the standard *Cell Ranger* reference transcriptome compared to the optimised transcriptome described by Pool et al. (2023) [10]. Pairs of light and dark versions of each colour represent the sample pairs, where samples with an 'optref' label refer to those processed using the optimised reference transcriptome. **(B)** Comparison of quality control metrics after quality control filtering of samples processed using the standard *Cell Ranger* reference transcriptome compared to optimised transcriptome described by Pool et al. (2023) [10]. Pairs of light and dark versions of each colour represent the sample pairs, where samples with an 'optref' label refer to those processed using the optimised reference transcriptome. **(C)** Comparison of *CellTypist* cell type annotation (using the *Pan_Fetal_Human* reference model) probabilities of samples processed using the standard *Cell Ranger* reference transcriptome compared to optimised transcriptome described by Pool et al. (2023) [10,13,14,16]. Annotation probabilities range between 0 and 1 and are binned in 10% increments. Probabilities for each sample are shown in separate plots and thus have differing y-axis scales due to the large differences in the number of cells recovered in each sample.

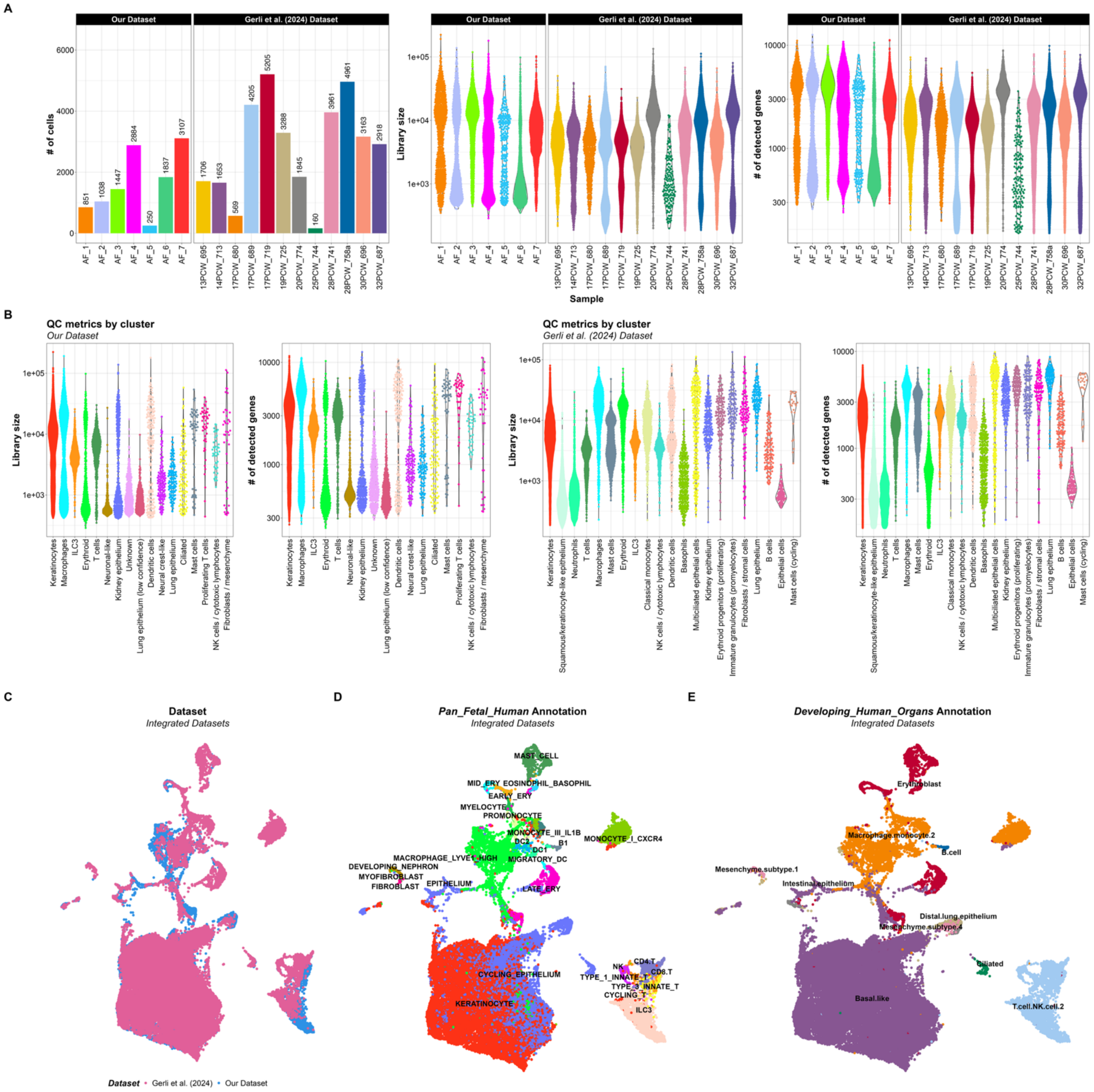


**Supplementary Figure 2. (A)** Comparison of quality control metrics by sample (number of cells per sample, library size and number of detected genes per cell) from our dataset and the Gerli et al. (2024) dataset after performing quality control filtering [1]. **(B)** Comparison of quality control metrics per cluster (library size and number of detected genes per cell) from our dataset and the Gerli et al. (2024) dataset after performing quality control filtering [1]. **(C)** UMAP plot of the integrated datasets coloured by dataset source. **(D)** UMAP plot of the integrated datasets coloured and labelled by

*Pan_Fetal_Human* annotations [13,14,16]. **(E)** UMAP plot of the integrated datasets coloured and labelled by *Developing_Human_Organs* cell type annotations [13–15].

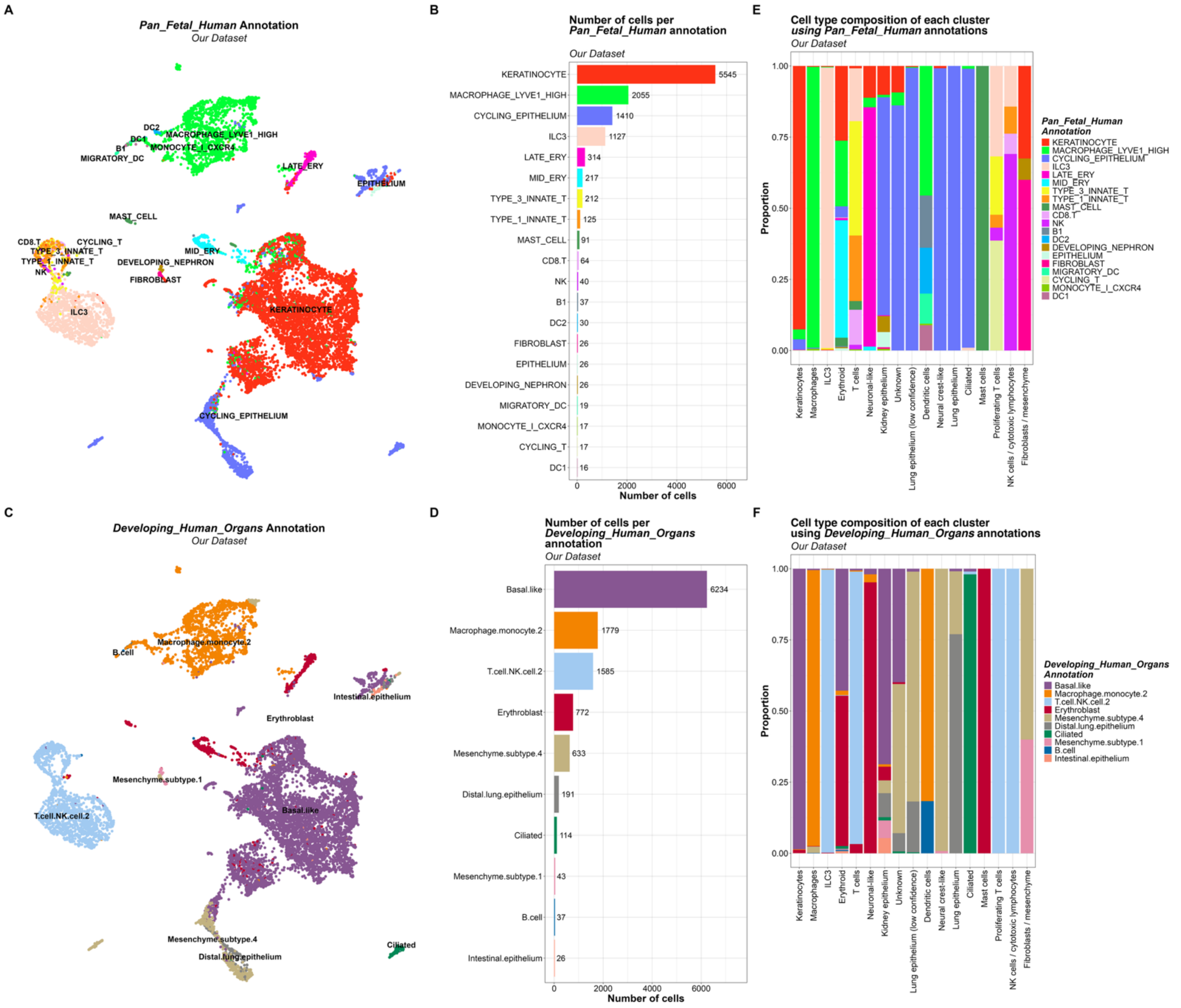


**Supplementary Figure 3.** ***(A)*** UMAP plot of our dataset, coloured by cell type annotation obtained from *CellTypist* using the *Pan_Fetal_Human* reference model [13,14,16]. **(B)** Number of cells per *Pan_Fetal_Human* cell type annotation in our dataset[13,14,16]. **(C)** UMAP plot of our dataset, coloured by cell type annotation obtained from *CellTypist* using the *Developing_Human_Organs* reference model [13–15]. **(D)** Number of cells per *Developing_Human_Organs* cell type annotation in our dataset [13–15]. **(E)** Cell type composition of each cluster in our dataset using *Pan_Fetal_Human* annotations. **(F)** Cell type composition of each cluster in our dataset using *Developing_Human_Organs* annotations.

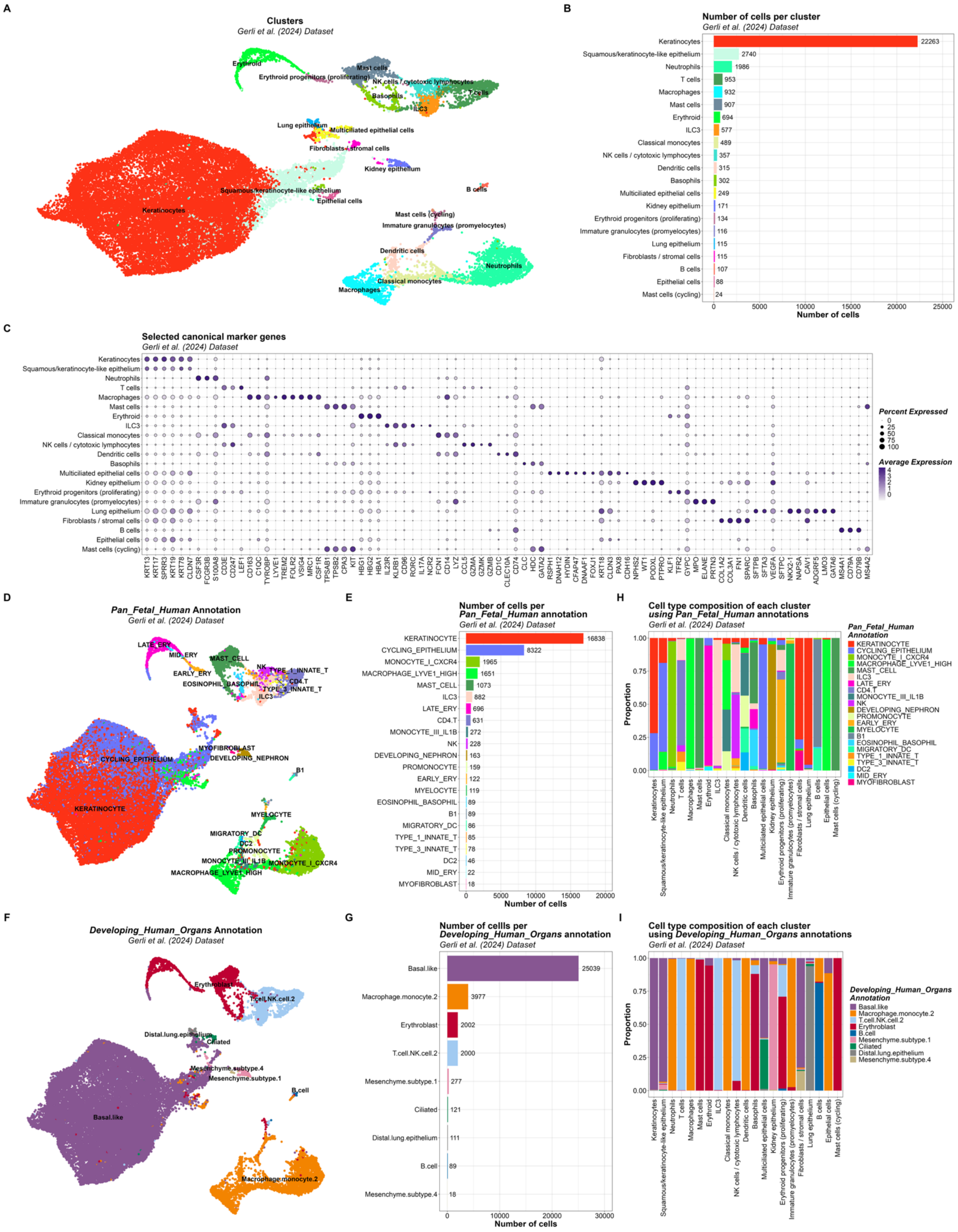


**Supplementary Figure 4.** **(A)** UMAP plot of the Gerli et al. (2024) dataset, coloured by cluster label [1]. **(B)** Number of cells per cluster label in the Gerli et al. (2024) dataset. **(C)** Dot plot of the expression of selected marker genes for each cluster in the Gerli et al. (2024) dataset. **(D)** UMAP plot of the Gerli et al. (2024) dataset, coloured by cell type annotation obtained from CellTypist using the *Pan_Fetal_Human* reference model [13,14,16]. **(E)** Number of cells per *Pan_Fetal_Human* cell type annotation in the Gerli et al. (2024) dataset. **(F)** UMAP plot of the Gerli et al. (2024) dataset, coloured by cell type annotation obtained from CellTypist using the *Developing_Human_Organs* reference model [13–

15]. **(G)** Number of cells per *Developing_Human_Organs* cell type annotation in the Gerli et al. (2024) dataset. **(H)** Cell type composition of each cluster in the Gerli et al. (2024) dataset using *Pan_Fetal_Human* annotations. **(I)** Cell type composition of each cluster in the Gerli et al. (2024) dataset using *Developing_Human_Organs* annotations.

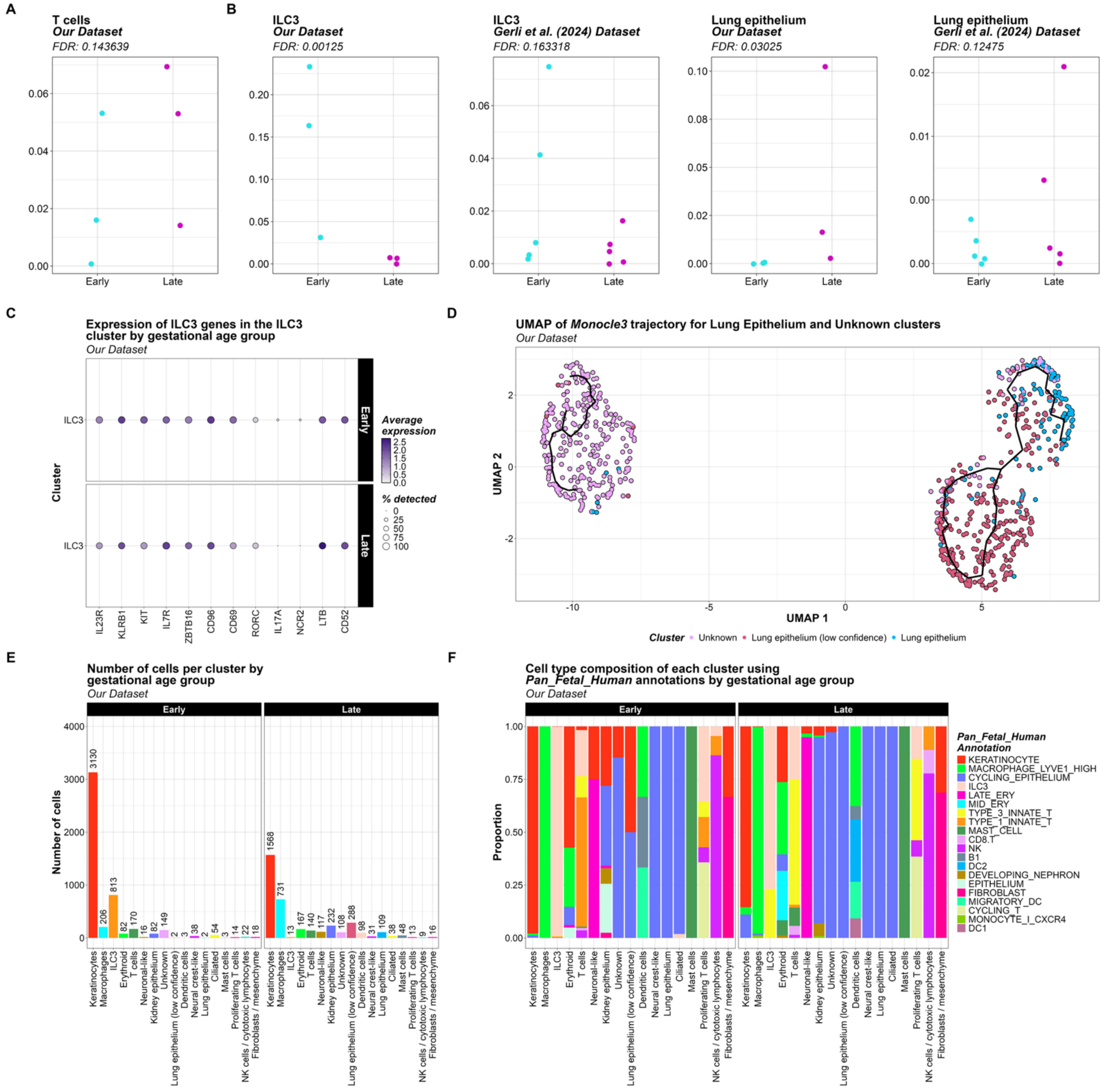


**Supplementary Figure 5. (A)** Dot plot showing the cluster proportion values for each sample in the early and late gestational age groups for the T cell cluster in our dataset. The FDR result of the *sccomp* test for each of these clusters is also shown above each plot [24]. **(B)** Dot plots showing the cluster proportion values for each sample in the early and late gestational age groups for the ILC3 and lung epithelium clusters in our dataset and the Gerli et al. (2024) dataset. The FDR results of the *sccomp* test for each of these clusters is also shown above each plot. **(C)** Dot plot showing the expression of *ILC3*-related genes in the ILC3 cluster for our dataset, split by gestational age group. Point colour reflects average expression value of that gene and point size reflects the percentage of cells it was detected in. **(D)** UMAP plot coloured by cluster, overlaid with the trajectory graph generated by *Monocle3* for the lung epithelium and unknown clusters in our dataset [25–27]. **(E)** Bar plots describing the number of cells per cluster in our dataset, split by gestational age group. **(F)** Bar plots describing the cell type composition of each cluster in our dataset, split by gestational age group. Cell type annotations were obtained from *CellTypist* using the *Pan_Fetal_Human* reference model [13,14,16].

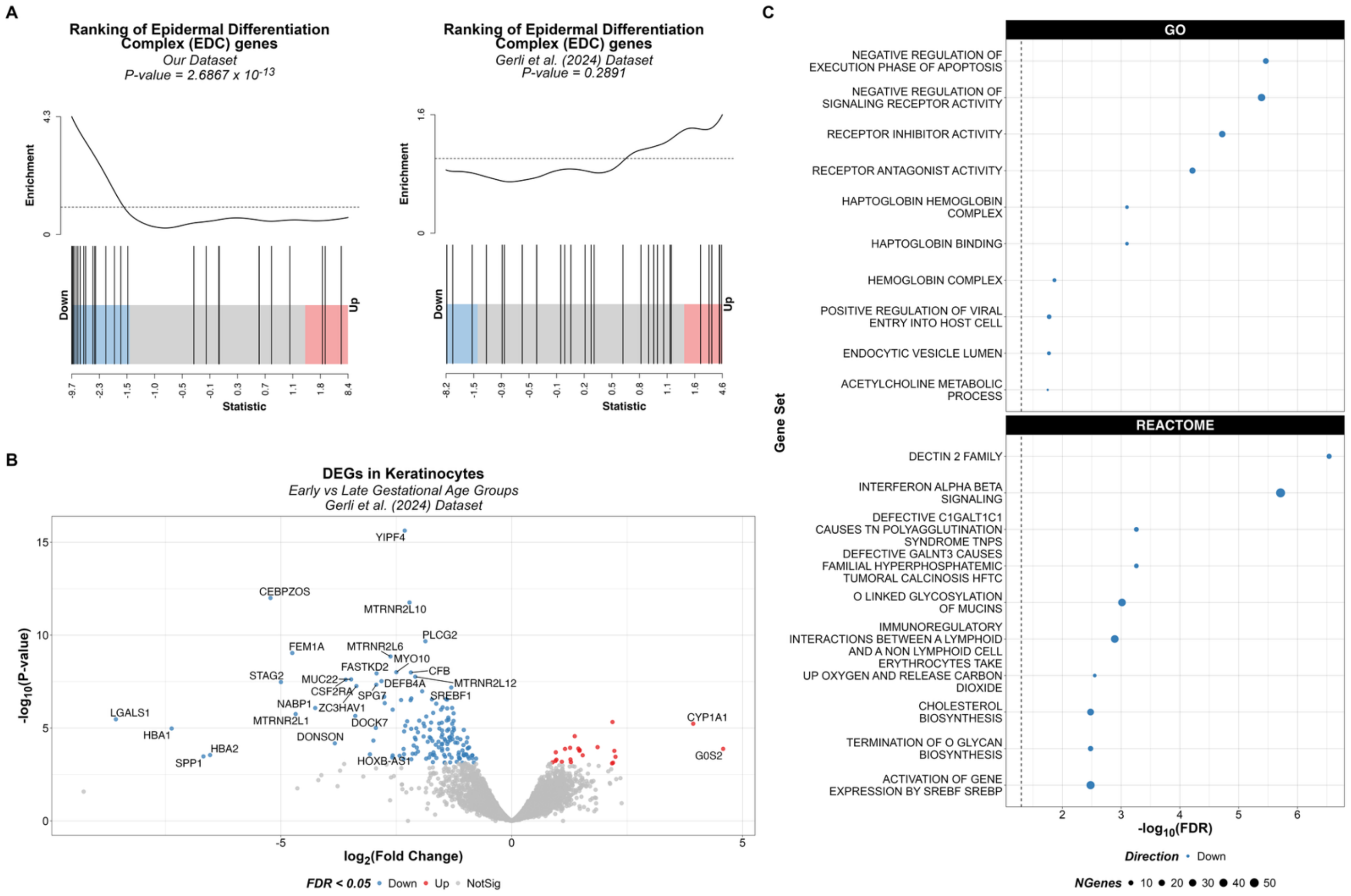


**Supplementary Figure 6. (A)** Barcode plots showing the enrichment of the EDC gene set in the DEGs for fetal skin cells (keratinocytes) between early (red; Up) and late (blue; Down) gestational age in our dataset and the Gerli et al. (2024) dataset. The p-value obtained from performing the *cameraPR()* competitive gene set test is displayed above each plot. **(B)** Volcano plot showing the DEGs in fetal skin cells (keratinocytes) between the early and late gestational age groups in the Gerli et al. (2024) dataset. Points are coloured by whether they were significantly upregulated in the late gestational age group (blue; Down), significantly upregulated in the early gestational age group (red; Up) or not significant (grey, NotSig). Points with an absolute log fold-change value above 3 or p-value below 0.0000001 were arbitrarily chosen to be labelled with their gene name. **(C)** Dot plots showing the GSEA enrichment results for DEGs identified for the Gerli et al. (2024) dataset using the GO and Reactome gene sets. Points are coloured by whether they were significantly upregulated in the late gestational age group (blue; Down) or significantly upregulated in the early gestational age group (red; Up; however, none of these are present). Point size reflects number of genes in the tested gene set.

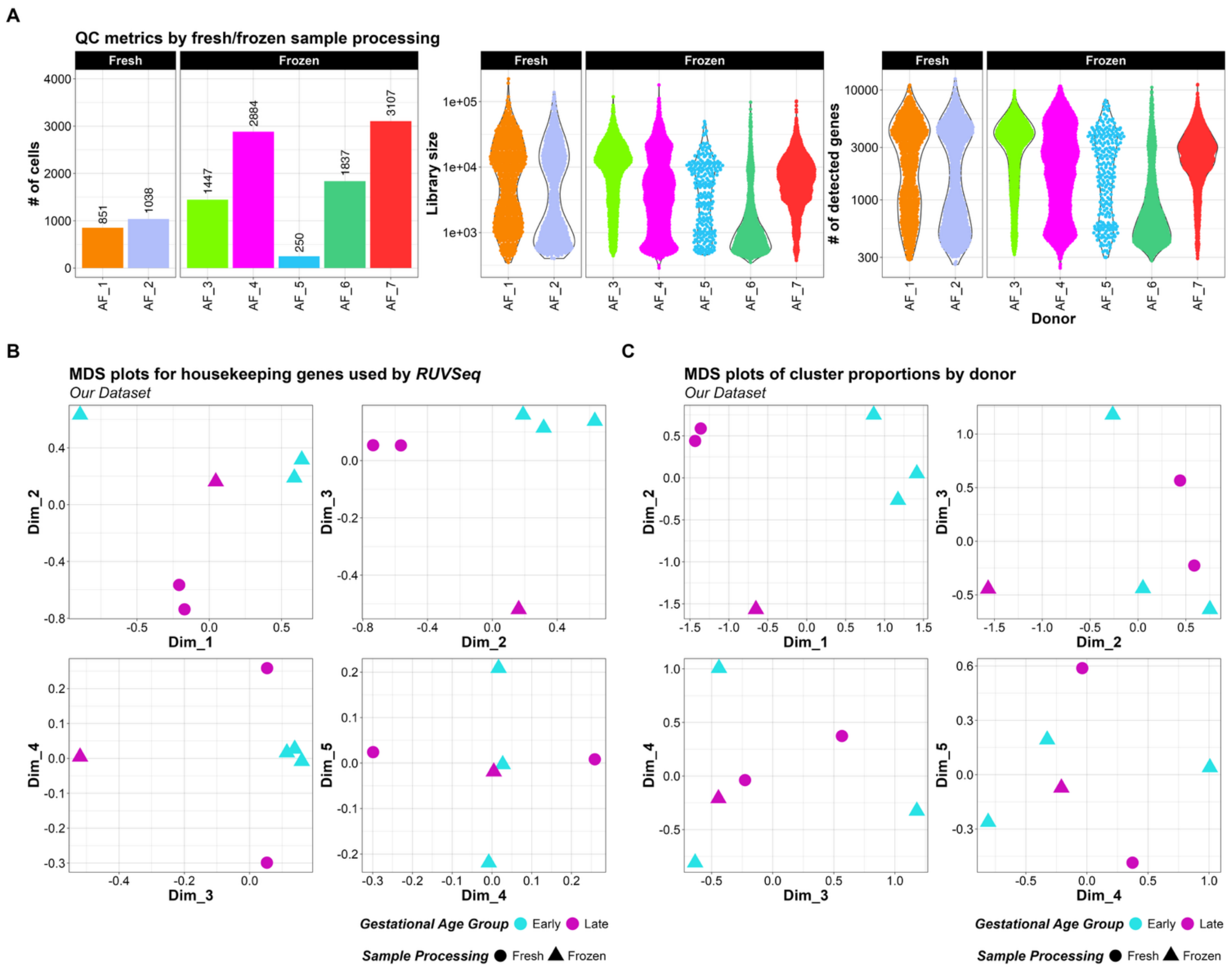


**Supplementary Figure 7. (A)** Comparison of quality control metrics per sample (number of cells per sample, library size and number of detected genes per cell) for our dataset, grouped by whether the sample was processed fresh or frozen. **(B)** MDS plots generated from count data from housekeeping genes for our dataset, pseudobulked by donor. Points (donors) are coloured by their gestational age group, and their shape reflects whether the sample was processed fresh or frozen. **(C)** MDS plot generated from cluster proportions for each donor. Points (donors) are coloured by their gestational age group, and their shape reflects whether the sample was processed fresh or frozen.
